# Cas9-induced nonhomologous recombination in *C. elegans*

**DOI:** 10.1101/2023.01.19.524763

**Authors:** Stefan Zdraljevic, Laura Walter-McNeill, Heriberto Marquez, Leonid Kruglyak

## Abstract

Identification of the genetic basis of phenotypic variation within species remains challenging. In species with low recombination rates, such as *Caenorhabditis elegans*, genomic regions linked to a phenotype of interest by genetic mapping studies are often large, making it difficult to identify the specific genes and DNA sequence variants that underlie phenotypic differences. Here, we introduce a method that enables researchers to induce targeted recombination in *C. elegans* with Cas9. We demonstrate that high rates of targeted recombination can be induced by Cas9 in a genomic region in which naturally occurring recombination events are exceedingly rare. We anticipate that Cas9-induced nonhomologous recombination (CINR) will greatly facilitate high-resolution genetic mapping in this species.

## Description

A central goal of modern genetics research is to identify genetic differences that influence phenotypic variation. However, studies that attempt to identify associations between genetic and phenotypic variation are often limited in resolution by the number and location of meiotic recombination events in the studied population. As a result, such studies often identify large genomic regions spanning many genes that harbor DNA sequence variants, and cannot further refine them to the underlying causal genes and variants. This is especially true for genetic mapping studies in *Caenorhabditis elegans*, a species characterized by a single meiotic recombination event per linkage group per generation and by extensive long-range linkage disequilibrium (Andersen et al., 2012; Barnes et al., 1995; Meneely et al., 2002; Rockman & Kruglyak, 2009). Therefore, new approaches to increase the rate of meiotic recombination are required to expedite the identification of causal genetic differences that underlie genomic regions associated with trait variation. Previous work in *Pristionchus pacificus* found that CRISPR/Cas9-induced double-stranded breaks can induce recombination (Lightfoot et al., 2019). We sought to determine whether we could facilitate genetic fine mapping in *C. elegans* by generating targeted recombination events with Cas9 in a similar manner.

To test our ability to target recombination, we first constructed a near-isogenic line (NIL) that contained a chromosome V region from strain XZ1516 introgressed into strain QX1211 (QX2501: qqIR51(V:21,547,828-22,058,188, XZ1516>QX1211)) (Figure 1A). We designed primers that amplify deletion variants in the QX1211 background at the 5’ and 3’ ends of the introgressed region to enable us to differentiate between the XZ1516 and QX1211 genotypes. The PCR band patterns for recombinant progeny are expected to be 5’ HET and 3’ HOM or 5’ HOM and 3’ HET, while double-recombinant progeny will be homozygous at both markers, with each parental genotype represented (Figure 1B). To establish a baseline recombination frequency within the introgressed region, we generated heterozygous F1 animals by crossing the NIL QX2501 with the QX1211 parent and genotyped 96 F2 self-progeny for recombination events. Of the 96 F2 progeny we attempted to genotype, we successfully amplified both PCR products for 76 individuals, in which we observed zero recombination events (Figure 1).

**Figure 1:**
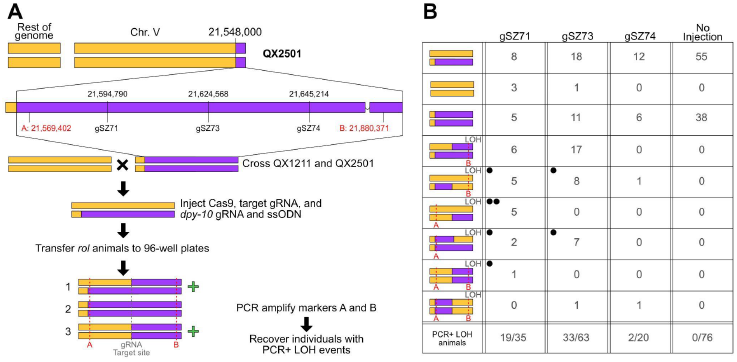
Cas9-induced nonhomologous recombination in *C. elegans*. A) Overview of the workflow used to induce nonhomologous recombination. A cartoon depiction of the genome composition of the QX2501 NIL used in this study is shown, where yellow represents QX1211 and purple represents XZ1516. Genomic coordinates (with XZ1516 as the reference genome) of the three gRNAs used in this study are shown below the genome representation of the NIL. Genomic coordinates of two indel variants used to differentiate between the parental genotypes are shown in red and labeled A and B. Young adult QX1211/QX2501 heterozygotes are injected with a gRNA that targets Cas9 to the desired location, along with a co-injection marker (*dpy-10* gRNA + ssODN repair template). The progeny of the injected animals that express the co-injection phenotype are singled to wells of a 96-well plate and allowed to self. A subset of the worms is genotyped with the A and B primer pairs. Individuals displaying loss of heterozygosity (LOH) such that both parental genotypes are still represented (i.e. individuals heterozygous at one marker and homozygous on the other, or homozygous at both markers) are marked with a green plus sign and recovered to agar plates for further characterization. B) Overview of the Cas9-induced nonhomologous recombination results. The table shows the PCR genotypes of *rol*/*dpy* progeny in this study. Left-most column depicts the expected strain genotypes based on PCR genotyping of the A and B indel variants. Within this column, recombinant genotypes are indicated by “LOH” in the upper-right corner. Dotted red lines indicate the marker where the LOH event occurred. The next three columns—one for each guide—show the number of genotyped *rol/dpy* progeny with the corresponding genotype. Black dots in the upper left corner correspond to the strains with whole-genome sequence data. The right-most column shows the F2 progeny genotypes when QX2501/QX1211 heterozygous animals were allowed to self without injection The last row of the table shows the total number of recombinants identified over the total number of F2 individuals genotyped for each gRNA.

Next, we designed three guide RNAs that span 96.5 kb of the 510 kb introgressed region to target double-stranded breaks. We generated Cas9 ribonucleoprotein injection mixtures for each of these guide RNAs and injected them into QX2501/QX1211 heterozygous animals (Paix et al., 2015). We transferred individual F2 progeny that exhibited the co-injection phenotype (*dpy*/*rol*) to 96-well plates and allowed them to self. Genotyping of the F3 progeny revealed high rates of recombination within the introgressed region for each of the three guides (gSZ71: 55.9% recombinants [19/35]; gSZ73 52.4% recombinants [33/63]; gSZ74: 10% recombinants [2/20]). While all three guide RNAs facilitated Cas9-induced recombination, we recovered only two recombinants using gSZ74, despite the fact that all three guides had comparable on-target scores (Doench et al., 2016). We hypothesize that the lower recombination induction efficiency of gSZ74 relative to gSZ71 and gSZ73 results from two QX1211 variants located in the gRNA target sequence. Given the high rates of Cas9-induced recombination we observed by PCR in a genomic region with undetectable natural recombination, we next sought to validate these findings via whole-genome sequencing of a subset of the recombinant strains.

We generated high-throughput sequence data for seven strains and found that all seven had Cas9-induced recombination events at the gRNA target sequence (Extended Data 1). Five of the sequenced strains had small insertion/deletion or single-nucleotide variants (SNVs) surrounding the edit site, while two had larger deletions (∼500 bp and ∼1500 bp). The strain with the fewest Cas9-induced differences from the reference XZ1516 genome and the QX1211 strain, QX2507, contained only a 3 bp insertion and a T>C SNV within the guide RNA target sequence. These results show that Cas9-induced double-stranded breaks can facilitate targeted recombination events at higher frequencies than natural recombination in *C. elegans*. Because new mutations were introduced near the gRNA target sequences, it is likely that the double-stranded breaks introduced by Cas9 are repaired by the non-homologous end joining (NHEJ) pathway. To minimize such introduced mutations during the generation of Cas9-induced nonhomologous recombination (CINR) events, we recommend PCR amplification and Sanger sequencing of the genomic regions surrounding gRNA target sequences in order to identify recombinants with the fewest Cas9-induced mutations, followed by whole-genome sequencing of these recombinants.

In this study, we show that Cas9-induced double-stranded DNA breaks facilitate targeted nonhomologous recombination in *C. elegans*. Notably, we were able to induce recombination at three distinct target sites in a genomic region with no detectable natural recombination. The approach presented here is an important advance for genetic fine-mapping because it will enable researchers to target maximally informative recombination events instead of having to rely on random recombination events. Because of the relative ease of generating targeted recombination events at a high frequency, we anticipate CINR will greatly expedite genetic fine-mapping in *C. elegans* in regions with low recombination rates, including chromosome centers (Rockman & Kruglyak, 2009) and divergent regions (Lee et al., 2021).

## Methods

### Strains

QX1211 and XZ1516 were acquired from the *Caenorhabditis elegans* Natural Diversity Resource (Cook et al., 2017). The *E. coli* strains OP50 and HB101 were acquired from the Caenorhabditis Genetics Center (CGC), which is funded by NIH Office of Research Infrastructure Programs (P40 OD010440).

### Worm maintenance

*C. elegans* strains were maintained at 20°C on modified nematode growth medium (NGMA), containing 1% agar and 0.7% agarose (Andersen et al., 2014) and seeded with OP50 *E. coli*.

### Crosses

QX1211 males were used to set up all crosses. For each cross, 5-10 hermaphrodites and 10-20 males were placed on 6 cm NGMA plates that were seeded with 10 µl of OP50 and allowed to mate overnight. The following day, individual plugged hermaphrodites were transferred to a new 6 cm NGMA plate and their progeny were monitored for a 50:50 male to hermaphrodite ratio.

### NIL construction

QX2501 was constructed from QX2500. We constructed QX2500 by generating F2 recombinants by selfing a QX1211/XZ1516 heterozygous strain and transferring them to individual wells of a 96-well plate with 50 µl of K media (Boyd et al., 2012) and 0.008 mg/ml HB101. Once the F2 strains generated progeny and the population starved, we transferred 15uL of the worm mixture to 15uL of 2X lysis buffer with 2% proteinase K and run on a thermocycler at 60 deg for one hour followed by 95 deg for 20 minutes. We used 2uL of this lysis mixture as template DNA for two PCRs using primer pairs: oZ50-51 and oZ52-53. These PCRs enabled us to identify recombinants within a defined genomic region. We recovered recombinants on 6 cm NGMA plates with OP50 and singled out 8 F3 progeny. We identified homozygous recombinant F3 progeny and backcrossed them to QX1211 for 6 generations, while maintaining the introgressed XZ1516 region. To generate QX2501, we injected young adult heterozygous QX2500/QX1211 animals with four target gRNAs (gSZ63, gSZ65, gSZ68, and gSZ70), the *dpy-10* gRNA and repair template (see Cas9 injections). We transferred F2 *rol* animals to 96-well plates, allowed them to self, and identified recombinant individuals using oZ66-67 and oZ64-65. We identified and isolated a recombinant that we named QX2501, which contains the QX1211 genotype from V:1-21,536,657, followed by a deletion spanning V:21,536,657-21,547,827, and the XZ1516 genotype from V:21,547,828-22,058,188.

### Cas9 injections

All guide RNAs were designed using the *multicrispr* R package (Bhagwat et al., 2020; Core, 2022). Guide RNAs were purchased from Synthego as synthetic spacer-scaffold fusions. Guide RNAs were resuspended in 30 µl of water and stored in −20ºC. Cas9 was purchased from IDT (cat #1081059) and stored in single-use aliquots (0.5 µl ∼ 0.5 µg Cas9) at −80ºC. Injection mixtures were made the same day as injections. For each injection mixture, a Cas9 aliquot was thawed on ice and mixed with 0.89 µl of the target gRNA (gSZ71, gSZ73, or gSZ74) and 0.89 µl of the *dpy-10* gRNA and incubated at 37ºC for 10 minutes in a thermocycler. After the incubation, 1 µl of the *dpy-10* single-stranded oligodeoxynucleotide (ssODN) repair template (oZ31), and 16.72 µl water was added to a final volume of 20 µl. The final concentrations in the injection mixtures were: 1.5 µM of Cas9, 4.45 µM of each gRNA, and 0.5 µM of the ssODN repair template.

### CINR Cas9 injections

The day before injections, L4 QX1211/QX2501 heterozygous animals were transferred to fresh 6 cm NGMA plates and allowed to develop overnight. The following day, young adults were injected and transferred to a fresh 6 cm NGMA plate to recover. The injected animals were singled to fresh 6 cm NGMA plates 16 hours after injections and monitored for the co-injection phenotype.

### CINR genotyping

The progeny of the injected animals that expressed the *rol* or *dpy* phenotypes (Arribere et al., 2014; Paix et al., 2015) were transferred to a 96-well assay plate (costar cat# 3370) and analyzed for recombination as described in the NIL construction section. We performed recombinant genotyping with oZ80-82-86 and oZ64-65 as separate PCR mixtures. PCR products were analyzed by gel electrophoresis to identify individuals displaying loss of heterozygosity (LOH) such that both parental genotypes are still represented (i.e. individuals heterozygous at one marker and homozygous on the other, or homozygous at both markers). A subset of LOH+ individuals were recovered to 6 cm NGMA plates and 8-16 F3 progeny were singled out and allowed to self. Once F4 progeny were present on the singled out F3 plates, the F3 parent was genotyped with the same PCR primer sets to identify individuals that were homozygous at the heterozygous marker in the F2 individual, such that both markers are homozygous with both parental genotypes represented.

### High-throughput sequencing

Genomic DNA was prepared using Purelink Genomic DNA Mini Kit (invitrogen cat# K1820-01), libraries were constructed using the Nextera XT library kit (illumina cat# FC-131-1024), and sequenced on the NextSeq500 (QX2500) or NextSeq2000 (QX2501-QX2508). Demultiplexing was performed on basespace. FASTQ files were aligned to the XZ1516 genome using *bwa mem* with default parameters (Li, 2013).

### Genome assembly scaffolding and annotation

We acquired the XZ1516 genome assembly from NCBI (project PRJNA692613) (Lee et al., 2021). We corrected misassemblies and scaffolding using the RagTag tool set (v2.0.1) (Alonge et al., 2021). To correct misassemblies, we used the following command: [*ragtag*.*py correct --mm2-param ‘-x asm20’*], using WS256 as the reference genome (Davis et al., 2022). To perform scaffolding, we used the following command: [*ragtag*.*py scaffold --mm2-param ‘-x asm20’*], using WS256 as the reference genome. We used funannotate (v1.8.8) (Palmer & Stajich, 2020) to perform gene annotation on the scaffolded assembly using the following command: [*funannotate predict -i XZ1516_ragtag_correct_scaffold_masked_nameChange*.*fasta -o funannotate_run_caenorhabditis -s “caenorhabditis” --strain XZ1516 --cpus 16 --organism other --busco_db nematoda --busco_seed_species nematoda --optimize_augustus*].

### Reagents

#### Strains

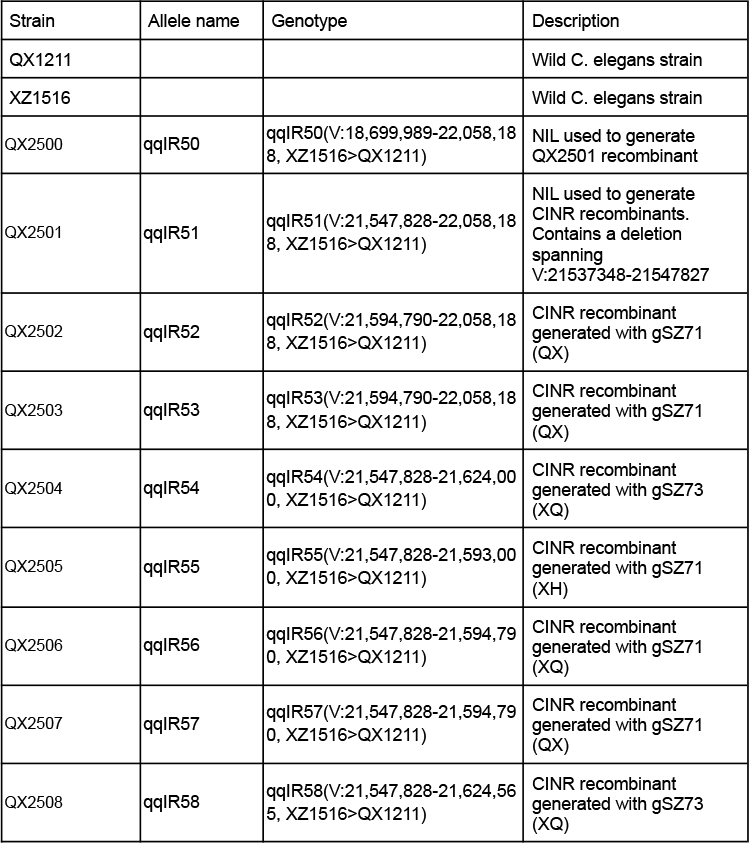

#### Oligos

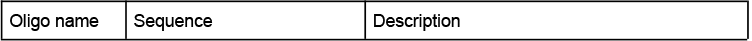
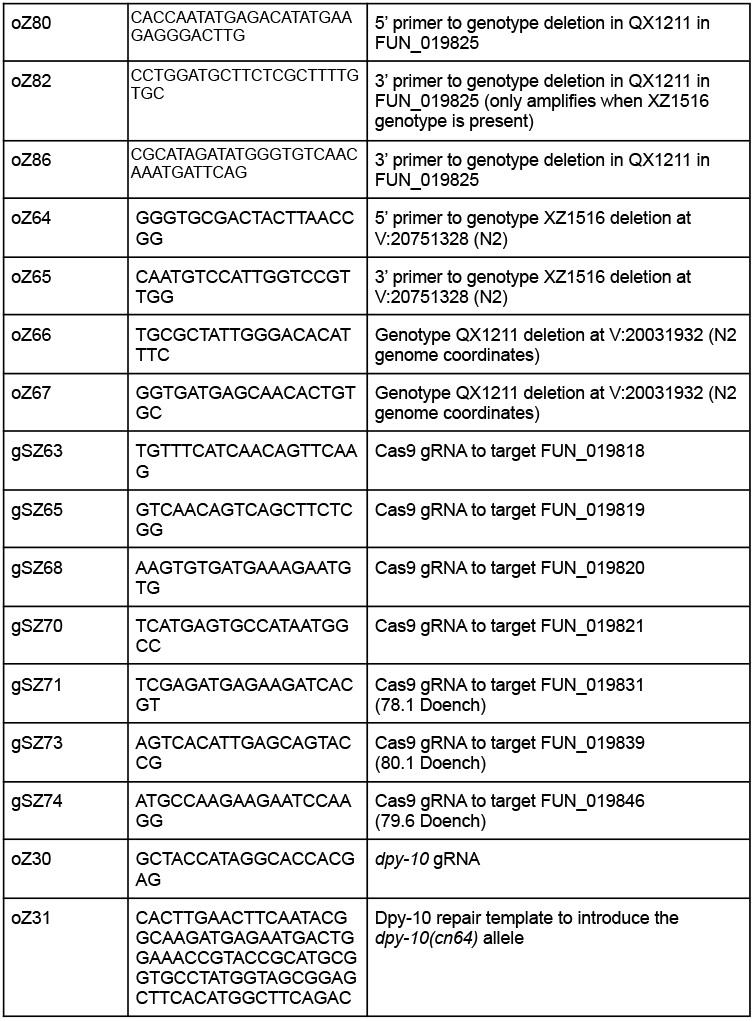
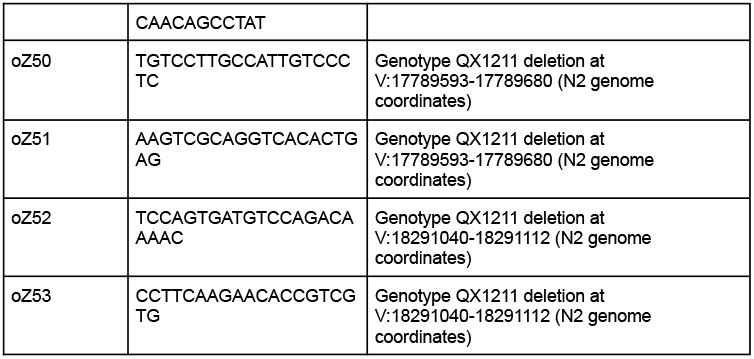

## Author contributions

Stefan Zdraljevic: Conceptualization, Data curation, Formal Analysis, Funding acquisition, Investigation, Methodology, Project administration, Supervision, Validation, Visualization, Writing – original draft, Writing – review & editing

Laura Walter-McNeill: Conceptualization, Data curation, Investigation, Methodology, Validation, Writing – review & editing

Heriberto Marquez: Investigation, Writing – review & editing

Leonid Kruglyak: Funding acquisition, Project administration, Resources, Supervision, Writing – review & editing

## Acknowledgments

We thank members of the Kruglyak lab for helpful discussions and for comments on the manuscript. We thank the CGC for bacterial strains, CeNDR for *C. elegans* strains, and the UCLA TCGB sequencing core for help with generation of high-throughput sequencing data.

