## Supplementary figures and images for "Cas9-induced nonhomologous recombination in *C. elegans*"

### Extended Data 1

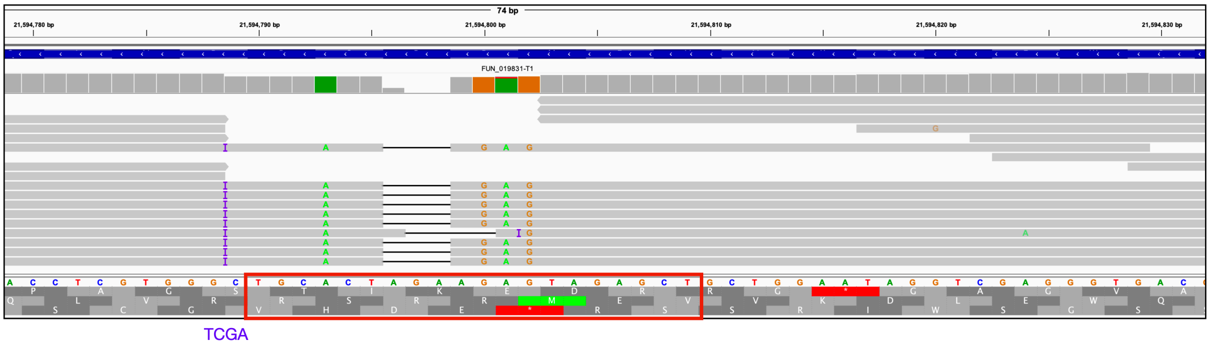

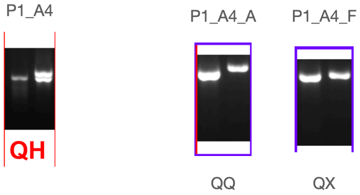

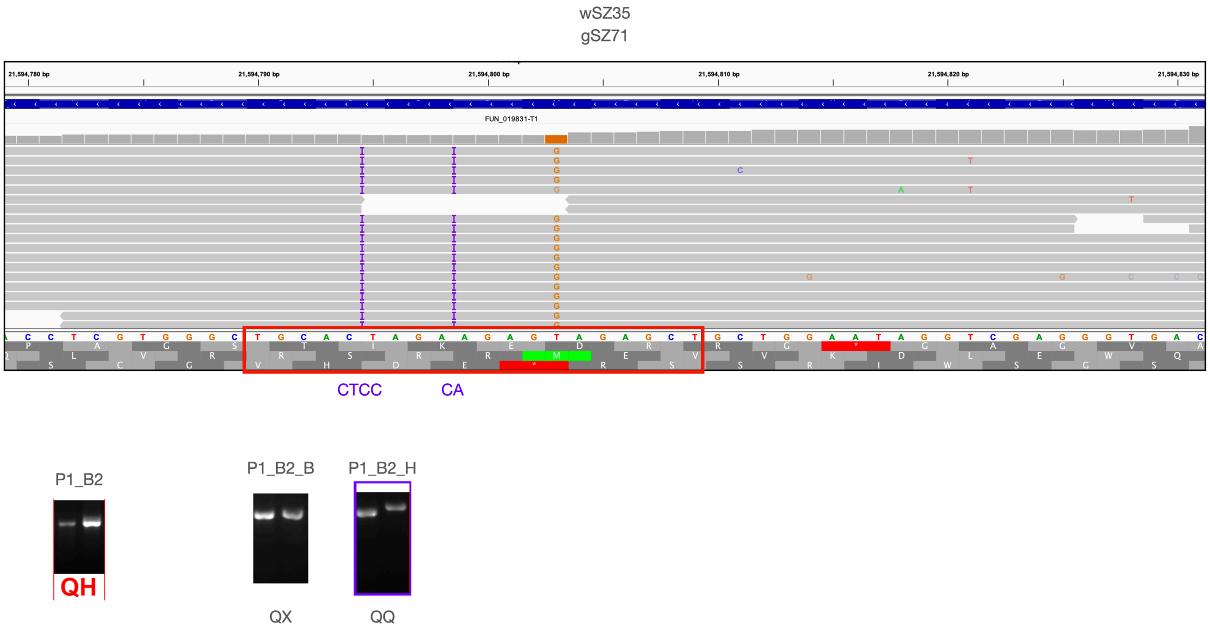

wSZ40
gSZ71

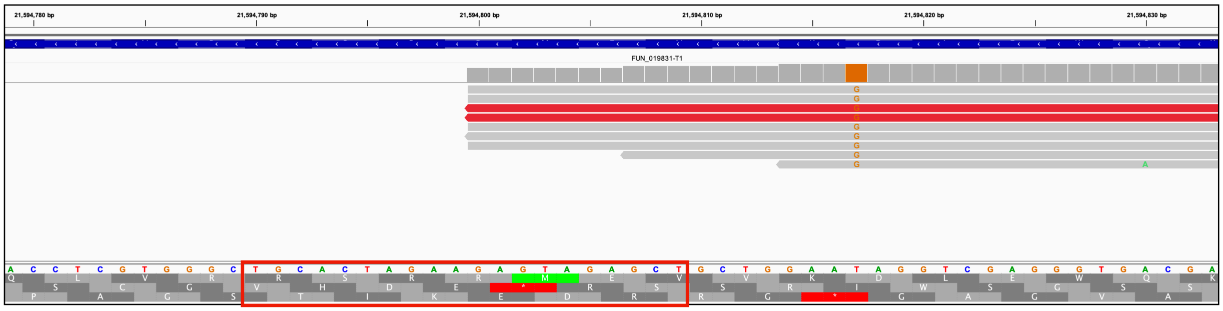

P1_C12 P1_C12_A

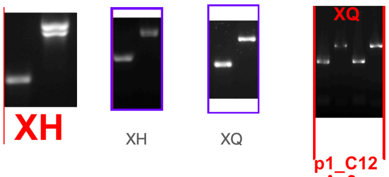

XH XH XQ p1_C12 _A_3

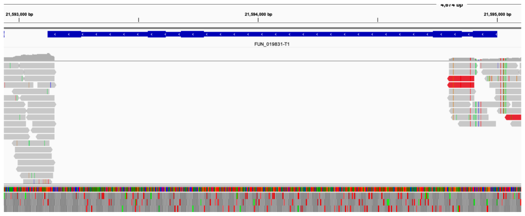

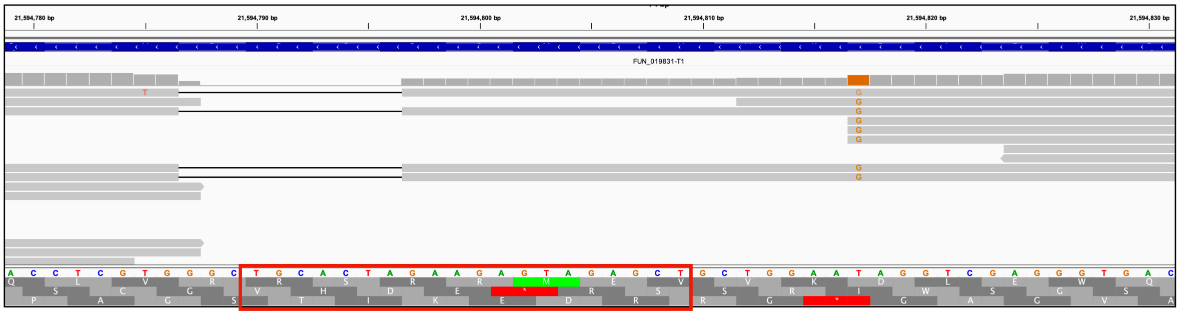

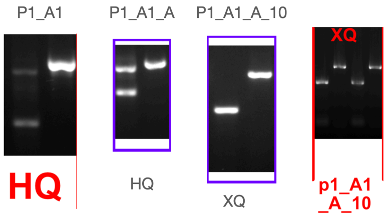

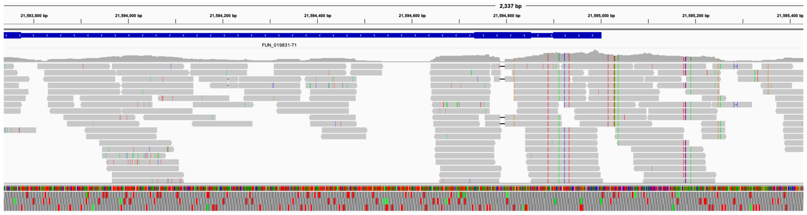

wSZ44

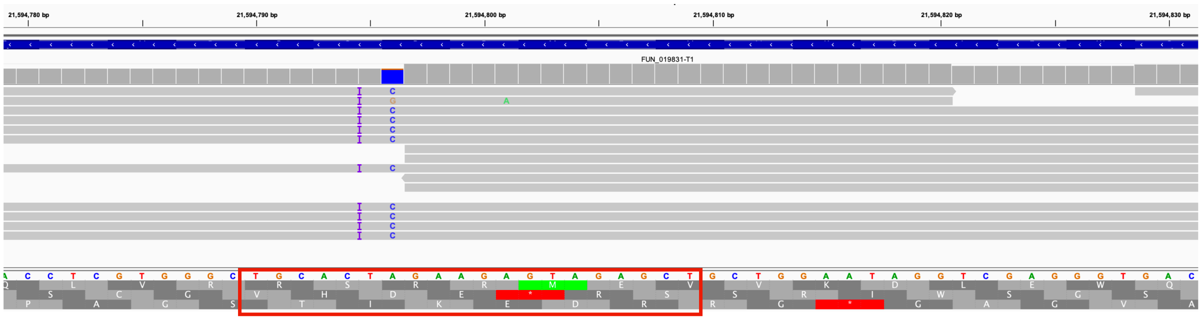


AGC

P1_A2

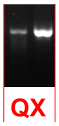


All QX

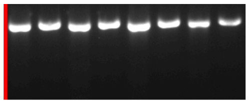

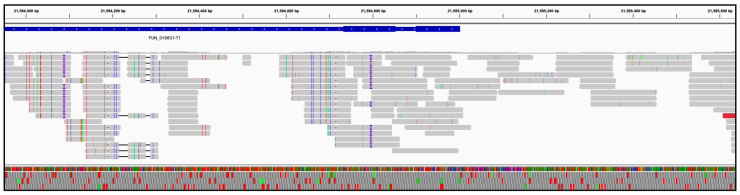

wSZ36

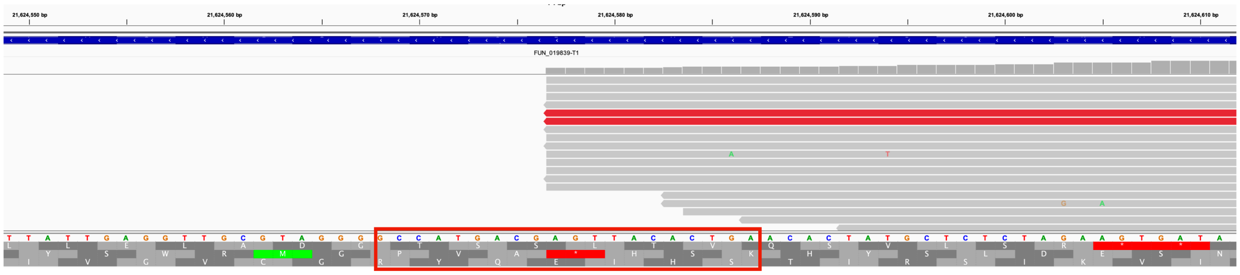

P1_F3

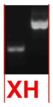


P1_F3_H

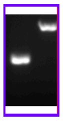

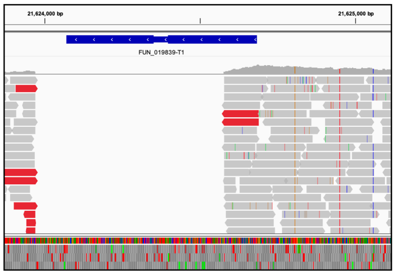

wSZ47

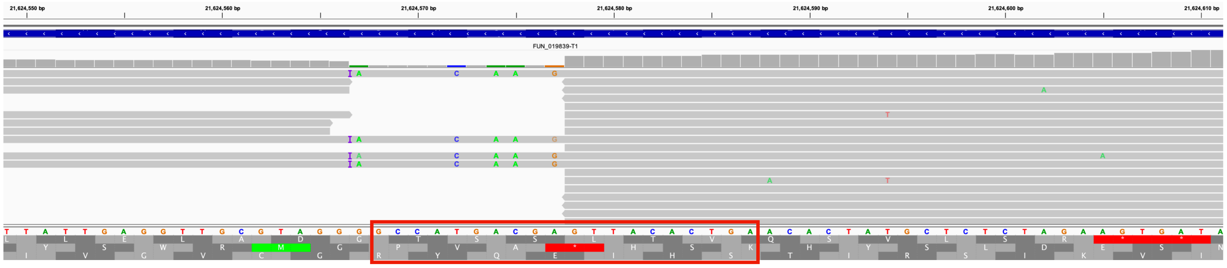

GTCACATTGGTCCA

P1_E7

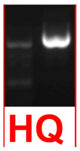


P1_E7_H

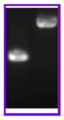

QX

All XQ

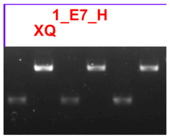


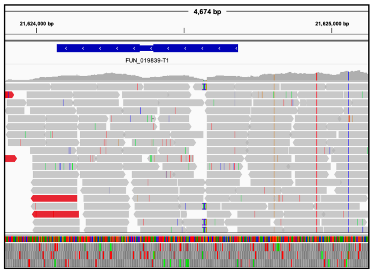

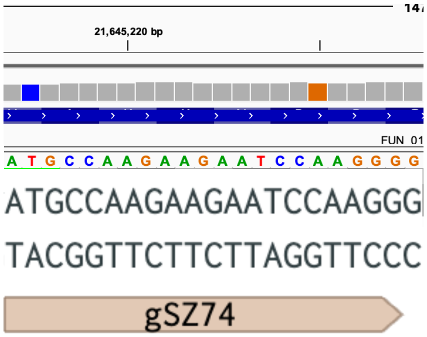

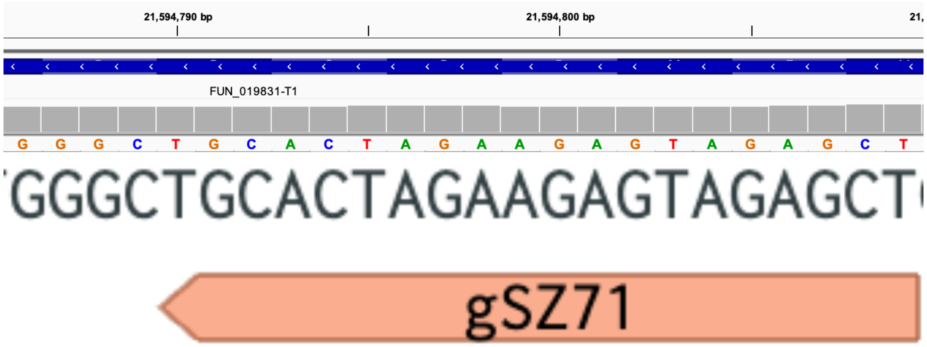

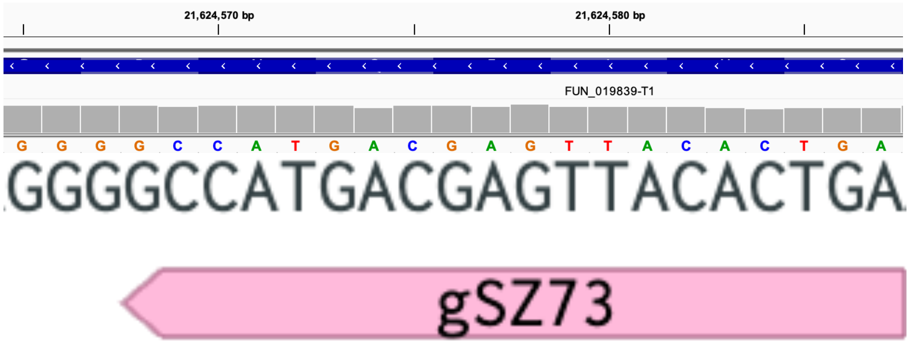
